# Chromosome-scale genome assembly and linkage map for *Silene uniflora* reveal the recombination landscape in a rapidly evolving plant species

**DOI:** 10.1101/2024.11.25.625183

**Authors:** Owen G Osborne, Daniel P Wood, Mariya P Dobreva, Luke T. Dunning, Rachel Tucker, Sarah ER Coates, Jaume Pellicer, Jon Holmberg, Adam C Algar, Greta Bocedi, Cecile Gubry-Rangin, Leonel Herrera-Alsina, Berry Juliandi, Lesley T Lancaster, Pascal Touzet, Justin MJ Travis, Alexander ST Papadopulos

## Abstract

The genus *Silene* is an important model system for fields as diverse as sex chromosome evolution, speciation and disease ecology. However, genomic resources remain scarce in the genus. Here, we present a chromosome-scale genome assembly for *S. uniflora*, a hermaphroditic/gynodioecious species which is an important model for rapid adaptation to anthropogenic disturbance and the role of phenotypic plasticity in adaptive evolution. Using a combination of long-read and Hi-C sequencing technologies, we generated a 1,268 Mb genome assembly with a scaffold N50 of 40.72 Mb and 682 Mb assembled into 12 chromosomes. We annotated the genome using evidence from transcriptome and protein mapping in combination with *ab initio* gene prediction, resulting in 41,603 protein-coding genes and a BUSCO completeness score of 91%. We also present a linkage map which we used to validate the genome assembly and estimate local recombination rate across the genome. Comparison to the only two other *Silene* species with chromosome-scale genome assemblies reveals widespread genome rearrangements in the genus, suggesting *Silene* may be a promising study system for the role of genome rearrangement in evolution, particularly in the evolution of sex chromosomes and adaptation.

**Significance statement:** Plant species in the genus *Silene* (campions) are important study organisms in multiple areas of ecology and evolution. Sea campion (*Silene uniflora*) is an important model for investigations into rapid adaptation, phenotypic plasticity and parallel evolution. However, only two species have high-quality genome assemblies available, hampering studies of their genetics and evolution. We present a high-quality genome assembly, genetic map and gene annotation for sea campion. These will be important genomic resources for future studies of sea campion, other species in the genus *Silene* and the family Caryophyllaceae more generally.

## Introduction

Sea campion, *Silene uniflora*, is a perennial herb with a native range in coastal regions of Northern and Western Europe (Davy, Alan J.M. M Baker, et al. 2024) Morphological and genetic studies of *S. uniflora* and its close relative *S. vulgaris* in the mid-20^th^ century were fundamental to the development of the field of experimental taxonomy (Davy, Alan J.M. M Baker, et al. 2024; Marsden-Jones & Turrill 1957). Subsequently, it became a model species for the study of castrating sexually transmitted fungal diseases (Chung et al. 2012; Abbate et al. 2018). It has also attracted attention for its rapid adaptation to the extreme environments resulting from non-ferrous metal mining (Baker & Dalby 1980; Baker 1978, 1974). More recently the species has become an emerging model system for research into the genetic basis of parallel evolution (Papadopulos et al. 2021) and the role of plasticity and gene expression during rapid adaptation (Wood et al. 2023; Coates et al. 2024). More broadly the genus *Silene* has been the focus of intense genetic research to understand sex chromosome evolution (Papadopulos et al. 2015; Yue et al. 2023), cytonuclear incompatibilities (Garraud et al. 2011; Postel et al. 2022), the evolution of mitochondrial genomes (Sloan et al. 2012; Wu et al. 2015), biological invasions (Keller & Taylor 2010; Jenkins & Keller 2011; Castillo et al. 2014), adaptation and speciation (Bratteler et al. 2006; Muir et al. 2012; Favre et al. 2017; Zemp et al. 2018). Although the value of genomic resources for the genus has been clear for some time, the first chromosome scale assembly was published only recently, prior to which assemblies were highly fragmented and incomplete (Fields et al. 2023).

There are currently only high-quality reference genomes available for two species of *Silene* - *Silene latifolia* and *S. conica* (Fields et al. 2023; Yue et al. 2023). *S. latifolia* is dioecious and has been the focus of intense research into the evolution of young, homomorphic sex chromosomes. The majority of *Silene* species are diploid with 12 pairs of chromosomes (2*n* = 24, including *S. latifolia* and *S. uniflora*; Bari 1973). *S. conica* is purely hermaphroditic and is unusual as it only possesses ten chromosome pairs (2*n* = 20; Fields et al. 2023). *S. latifolia* and *S. conica* also sit at the extreme ends of genome size for *Silene* species – *S. latifolia* has one of the largest diploid genome sizes (2.8 Gb; Pellicer & Leitch 2020), whereas *S. conica* is one of the smallest (0.9 Gb; Williams et al. 2021). Populations of *S. uniflora* can exclusively contain hermaphrodites or be gynodioecious (Davy, Alan J.M. Baker, et al. 2024). Its genome size is intermediate for the genus (1.2 Gb), it has twelve pairs of chromosomes (Williams et al. 2021) and B chromosomes have also been observed in karyotyping studies (Cobon & Murray 1983). High quality genomic resources for the species have the potential to accelerate research into the genomic and epigenetic mechanisms that underly rapid adaptation, but also contribute to wider research in the genus by adding additional resources for studies of sex chromosomes and mating systems. Here, we report a chromosome-level genome assembly and a detailed genetic map for *S. uniflora*, providing a comprehensive resource for future eco-evolutionary research.

## Materials and methods

### Plant material and sequence data generation

Cuttings from a single coastal individual were collected in Tresaith (West Wales, UK) propagated and self-fertilized as part of a previous study (Papadopulos et al. 2021). A single individual inbred F1 (SUTF1P; draft genome ASM1898310v1) was selfed and the F2 individual sequenced here was grown under controlled conditions at the Henfaes Research Centre, Bangor University, UK. High molecular weight DNA was extracted for Pacific Biosciences (PacBio) and Oxford Nanopore (ONT) sequencing using a Qiagen DNAeasy Maxi plant kit. PacBio library preparation and sequencing was completed at the Genomics Laboratory, University of Sheffield, UK (Supplementary note 1). Liquid nitrogen frozen leaf tissue was sent to Dovetail Genomics, LLC for Hi-C, Chicago and Illumina Truseq library preparation and sequencing on a HiSeqX sequencer (Illumina).

RNA was extracted from liquid nitrogen frozen leaves, flowers and roots separately using Qiagen RNeasy Plant Mini Kit, including the optional DNase digestion step. RNA extraction was conducted on root, flower and leaf tissue for the genome plant using the RNeasy Plant Mini Kit (Qiagen), including a DNase digestion step, and RNA extracts were sent to the Beijing Genomics Institute (Hong Kong) for library preparation and sequencing. 100 bp paired end RNA-seq libraries were prepared according to the BGISEQ-500 RNA-Seq Library Preparation Protocol and libraries were sequenced on a BGISEQ500 sequencer (Beijing Genomics Institute).

For linkage mapping, two plants were grown from seed collected from populations i) WW-C and ii) WW-M (Papadopulos et al. 2021) and were subject to a controlled cross. One of the offspring of this cross was selfed and the resulting seeds (F2s) collected (Supplementary note 2). When the plants were eight months old, leaf punches were taken for DNA extraction and sequencing using LGC Genomics GmbH SeqSNP service. LGC Genomics constructed SeqSNP sequencing libraries for F2 progeny using 25,000 custom probes (Table S1). These libraries were sequenced on an Illumina NextSeq 500.

### Genome size estimation

The nuclear DNA content of *Silene uniflora* was determined by flow cytometry following the one-step procedure described in Doležel et al. (2007; Supplementary note 3).

### Genome assembly

An initial draft hybrid assembly was constructed from raw Illumina short read data and PacBio longread data using the MaSuRCA pipeline (kmer size = 99, PacBio coverage = 30x). Contigs from this draft assembly were then arranged into scaffolds using the Hi-C and Chicago data using the Dovetail HiRise pipeline. To further improve the assembly, we utilised *Oxford Nanopore MinION* (ONT) long reads. ONT data were first corrected using the short reads produced from the Hi-C scaffolding (Supplementary note 4). Corrected long reads were first used to perform additional scaffolding of the assembly before being used to close assembly gaps. Scaffolding was conducted using SLR v1.0.0 (Luo et al. 2019) with a minimum alignment length of 300 nucleotides. Gap closing was conducted using *TGS-GapCloser* (v0.56; Xu et al. 2019). Contamination was assessed using the NCBI Foreign Contamination Screen pipeline (Astashyn et al. 2024) and any adaptor and non-target organism sequences were masked from the final assembly. For each step in assembly improvement, QUAST (Gurevich et al. 2013) was used to assess assembly size and contiguity.

### Genome annotation

Prior to gene prediction, we masked repeat regions using *RepeatMasker* (v4.1.0; http://www.repeatmasker.org/) against the MIPS plant repeat database (v.9.3; Nussbaumer et al. 2013). For gene annotation, we employed RNA-seq mapping, assembled transcript mapping, protein mapping and *ab initio* approaches, which were then combined into a single non-redundant gene model (Supplementary note 5). We used eggnog-mapper (v 2.0.1; database v2.0; Cantalapiedra et al. 2021) to functionally annotate predicted proteins. We used diamond as a mapper, a minimum query and subject coverage of 70%, an e-value cutoff of 0.0001, a taxonomic scope of Viridiplantae and defaults for all other options. To produce a final set of high-quality gene annotations, we filtered out annotations with non-canonical splice sites or in-frame stop codons, those annotated as transposable elements in eggnog-mapper, those without start or terminal stop codons, and those without either RNA evidence or matches to know ortholog groups in eggnog-mapper. Annotation completeness was assessed using BUSCO (v4; Simão et al. 2015) in protein mode with the Viridiplantae, Embryophyta and Eudicot gene sets.

### Linkage map construction and analysis

Raw reads from the SeqSNP data were trimmed and mapped to the genome (Supplementary note 6). Linkage maps were constructed using LepMap3 v0.4 (Rastas 2017; Supplementary note 7). The linkage map was used to provide validation of the genome assembly, indicate the position of unassigned scaffolds and estimate local recombination rate. Recombination rate was estimated using a LOESS local regression (span 0.1 Mb), with recombination rate estimated as the slope in non-overlapping 1 Mb windows. To avoid inaccurate recombination rate estimates caused by error or rearrangements between the plants used for genome assembly and linkage mapping, we did not calculate recombination rate for windows containing no markers or where the map window contained markers from other scaffolds.

### Comparative genomics

Synteny between the *S. uniflora* genome and the previously published assemblies of *S. conica* and *S. latifolia* was examined using the MCscan algorithm implemented in the JCVI toolkit v.1.3.8 (Wang et al. 2012; Tang et al. 2024; Supplementary note 8).

## Results and Discussion

### Genome assembly, linkage mapping and annotation

The mean estimated genome size from our flow-cytometry analysis was 1C = 1.28 pg, corresponding to a haploid genome size of 1251.48 Mb (Table S2). The final genome assembly was 1269.36 Mb long, closely matching the estimated genome size. The assembly contained 12,573 scaffolds, had a scaffold N50 of 40.72 Mb and a scaffold L50 of 11. Approximately 53.73 % of the assembly was contained within the 12 largest scaffolds which were between 32.17 and 76.86 Mb in size (henceforth chromosomes Chr01 – Chr12), corresponding to the haploid chromosome number of *S. uniflora*. Additionally, five further scaffolds were over 1 Mb in length. Contamination screening identified no foreign organism sequences and 5 putative adaptor sequences (totalling 134 bp; Table S3) which were masked from subsequent analysis. Starting from an initial assembly with an N50 of 34 Kb, each step in the genome assembly significantly improved genome contiguity (N50 values: Chicago: 169 Kb; Dovetail Hi-C: 27.01 Mb; ONT scaffolding: 34.97 Mb, ONT gap-filling: 40.72 Mb; Table S4). The final gene annotation contained 41,603 protein coding genes encoding 48,596 putative transcripts. Of these, 98.89% were complete, including both start and stop codons and 32.26% also contained annotations for 5’ and 3’ untranslated regions (Fig. S1). BUSCO analysis suggested that the annotation was largely complete (86% - 91% BUSCO completeness depending on the gene set used; Fig. S2).

The linkage map contained 8,372 markers on 1,357 scaffolds which were assigned to 12 linkage groups (LGs). The linkage map was used to validate the assembly of the chromosomes and estimate the correct placement of other large scaffolds. LGs were broadly concordant with assembled chromosomes. Between 87.2 and 97.5% (mean = 95.6) of markers from each chromosome were assigned to a single LG, and the LG differed for each chromosome (Table S5). The map suggested that three of the five unplaced scaffolds >1 Mb were linked to Chr12 in LG., one to Chr06, and one to Chr02 (Figs. S3-5). Some inconsistencies between the map and assembly may represent rearrangements between the populations that the genome and mapping plants originated from. For example, plots of physical versus map position highlight a putative intrachromosomal translocation on chromosome Chr07, and putative inversions on Chr01, Chr03, Chr06, Chr09 and Chr10 (Fig. S6). Frequent chromosomal rearrangements may have important implications in *S. uniflora*, since these have the potential to contribute to reproductive isolation between locally adapted populations (Kirkpatrick & Barton 2006).

Mean local recombination rate varied between chromosomes (ranging from 1.35 - 2.12 cM/Mb; Fig. S7) and was negatively correlated with chromosome length (Spearman’s rank correlation: *ρ* = -0.87; *P* = 0.0004; Fig. S8), a common finding across eukaryote species (Haenel et al. 2018). Within chromosomes, and on a genome-wide scale, local recombination rate was significantly positively associated with gene density, but not GC content, despite gene density and GC content being correlated with each other (Fig. 1, Fig. S9).

**Figure 1.**
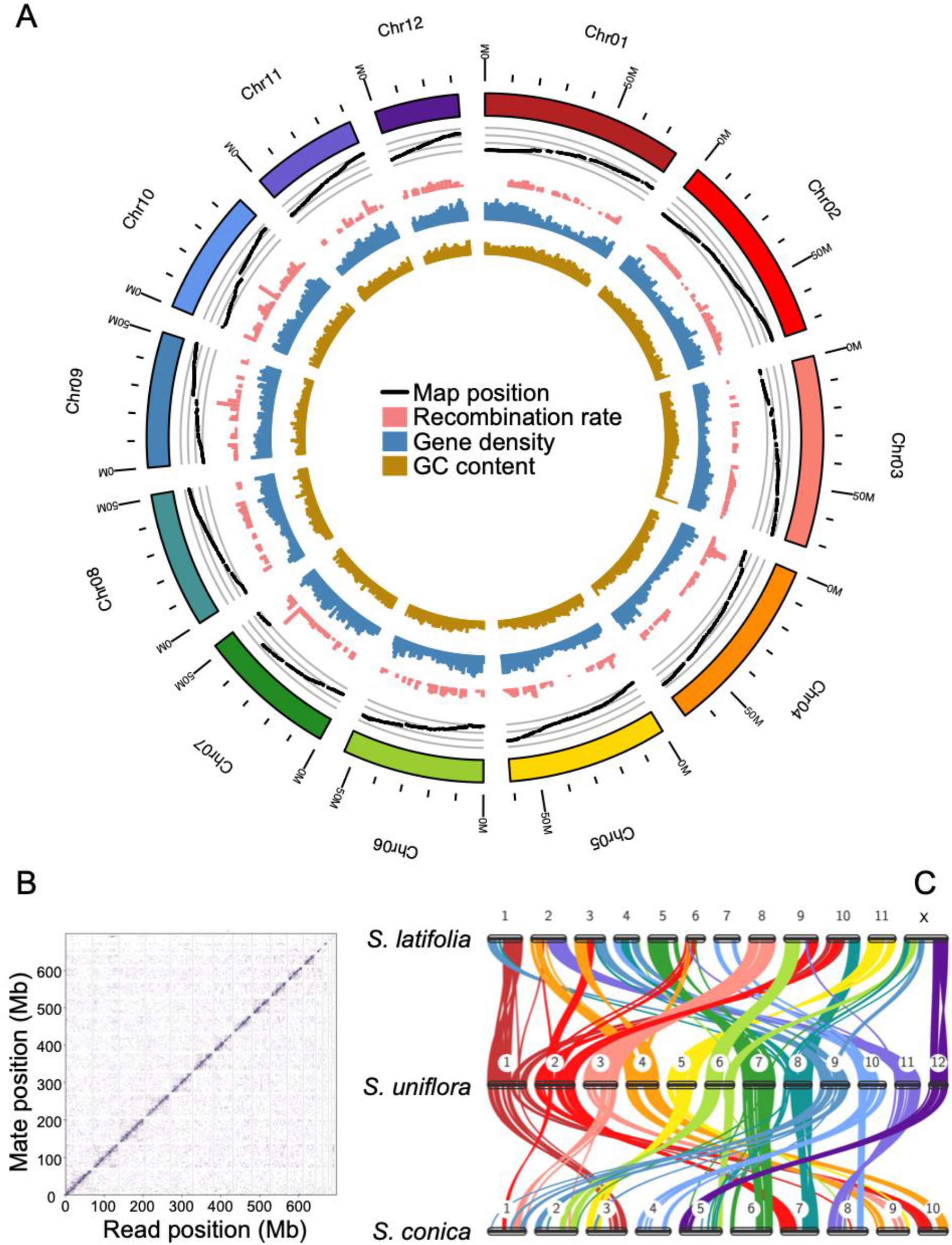
Genome assembly of *Silene uniflora* and comparison to other *Silene* genomes. The *S. uniflora* chromosomes are shown as a circular plot (A) with GC content (%), gene density (genes per 1Mb), and estimated recombination rate (cM/Mb) shown in non-overlapping 1Mb windows. The relationship between physical and genetic distance is shown as a “Marey map” (Chakravarti 1991), with each point representing one genome-anchored marker and the y-axis representing its position in our linkage map. The Hi-C scaffolding is shown as a link density histogram (B) with the genomic position of each read and its mate shown on the x and y axes respectively and colour intensity representing number of read-pairs per bin. Comparison of the *S. uniflora* genome with previously published *Silene* genomes shows widespread rearrangements between the species (C). Links between the chromosomes of each species represent syntenic blocks shared between the species, coloured by *S. uniflora* chromosome as in panel A.

Comparison to the *S. latifolia* and *S. conica* genomes revealed widespread rearrangements between the species. Notably, the X chromosome of *S. latifolia* was linked to four *S. uniflora* chromosomes. It has previously been suggested that chromosomal translocations have played a causative role in sex chromosome formation in *S. latifolia* and its sister species *S. dioica* (Bačovský et al. 2020). However, our synteny analysis suggests that most large-scale rearrangements in the *S. latifolia* X chromosome relative to *S. uniflora* are shared by *S. conica* – a hermaphrodite lacking sex chromosomes – suggesting these rearrangements occurred prior to the evolution of dioecy (Fig. 1C).

Here, we present a high-quality genome assembly and linkage map for *S. uniflora*, which we use to infer the recombination landscape of *S. uniflora* and investigate genome rearrangements between *Silene* species. This will be an important resource for studies using *Silene* as a model organism. *S. uniflora* populations have repeatedly and rapidly evolved tolerance to heavy metal contamination. A high-quality genome assembly will allow the role of genomic rearrangements in this process to be examined. Furthermore, as one of only three *Silene* species with a high-quality genome assembly, this resource is likely to be important for studies of other species and Caryophyllaceae more broadly.

## Author Contributions

ASTP, DPW, LD, RT and MPD produced the sequence data. DPW produced the linkage map with assistance from JH. OGO and ASTP conducted genome assembly, scaffolding, annotation and subsequent analysis with input from DPW, PT and SERC. ASTP, ACA, GB, CGR, BJ, LTL, JMJT supervised the project. OGO wrote the first version of the manuscript, and all authors contributed to and approved the final version of the manuscript.

## Acknowledgements

The work was funded by a UK Natural Environment Research Council grant to ASTP (NE/R001081/1). We thank, Daniel Sloan and Peter Fields for helpful discussions on the analysis, Nick Welsby for laboratory support, and Ade Fewings and Supercomputing Wales for computational support and resources.

