## Supplementary information for "Chromosome-scale genome assembly and linkage map for *Silene uniflora* reveal the recombination landscape in a rapidly evolving plant species"

**Supplementary note 1: long-read sequencing details.**
A SMRTbell library was constructed using the Sequel Sequencing Kit 2.1 v2.0 (Pacific Biosciences, Menlo Park, CA). The SMRTbell library was sequenced using four Sequel™ SMRT® Cell 1M v2 Tray, on the PacBio Sequel system with a collection time of 10 hours. For Oxford Nanopore (ONT) sequencing, the Circulomics Short Read Eliminator XL Kit (Pacific Biosciences) was used to enrich DNA for high molecular weight fragments, and the Genomic DNA by Ligation Sequencing Kit (LSK109, Oxford Nanopore Technologies), the NEBNext Companion Module for ONT Ligation Sequencing (New England Biolabs), and Agencourt AMPure XP beads (Beckman Coulter Inc) were used to prepare the sequencing library. ONT sequencing was conducted on a single R9.4 flow cell on a MinION v2.

**Supplementary note 2: Seedling growth conditions.**
The seedlings were grown in Erin Traditional 24 Multipurpose Compost in 80ml cells in a greenhouse (16/8 hr day/night cycle; temperature controlled 18/12 ºC, supplementary lighting automatically switched on if light levels fell below 120 µmol $m^{-2}$ $s^{-1}$ during the day) before being transplanted to 1.5 L pots filled with compost and 14.7g MiracleGro Slow Release Fertiliser after five weeks.

**Supplementary note 3: Genome size estimation.**
The nuclear DNA content of *Silene uniflora* was determined by flow cytometry following the one-step procedure described in Doležel et al. (2007). Briefly, ~1 $cm^2$ of fresh leaf material of *S. uniflora* and the calibration standard (*Pisum sativum* 'Ctirad', 2C = 9.09 pg; Doležel et al. 1998) were chopped together in 2 ml of nuclei isolation buffer (Ebihara et al. 2005), filtered through a 100 µm Celltric filter (Sysmex), and stained by adding 100 µl of 1 mg $ml^{-1}$ propidium iodide solution. Nuclei suspensions were then incubated for ~20 min on ice prior to analysis. The relative fluorescence of stained nuclei was acquired using a Cyflow Space (Sysmex-Partec GmbH, Münster, Germany) flow cytometer fitted with a 100 mW green (532 nm) solid-state Cobalt Samba laser. The resulting flow histograms were analysed using the software FloMax v.2.4. Three leaves from three *S. uniflora* individuals held in the living collection of the Royal Botanic Gardens, Kew were measured separately, and a minimum of 1,000 nuclei per fluorescence peak were recorded in each analysis.

**Supplementary note 4: ONT read correction.**
Short reads were first processed to remove adaptor sequences and low quality bases using *Cutadapt* (version 2.3; Martin 2011) and *Trimmomatic* (v0.38; using the following settings: TRAILING:10, SLIDINGWINDOW:4:15, MINLEN: 64; Bolger et al. 2014). A Burrows-Wheeler Transform was constructed from the trimmed *Illumina* reads using *ropebwt2* (Li 2014) which was then used to correct long-reads using *FMLRC* (version 1.0.0; Wang et al. 2018).

**Supplementary note 5: Genome annotation details.**
RNA-seq data was initially pre-processed to remove low-quality reads and sequencing adaptors using Trimmomatic (Bolger et al. 2014; v.0.39; with the following settings: LEADING:10 82 TRAILING:10 SLIDINGWINDOW:4:15 MINLEN:70). For the RNA-seq mapping approach, cleaned RNA-seq reads for all three tissues were mapped to the genome assembly using HISAT2 (v.2.1.0; Kim et al. 2019) using the "dta" option to produce output tailored for transcript assemblers. The resulting alignment was sorted using samtools (v.1.9; Li et al. 2009) and used to

produce gene predictions with StringTie (v.1.3.6; Pertea et al. 2015). For the assembled transcript mapping approach, cleaned RNA-seq reads were first used to produce a *de novo* transcriptome assembly using Trinity (version 2.10.0) using default settings (Haas et al. 2014). Assembled transcripts were mapped to the genome using PASA v2.4.1 (Haas et al. 2008) using GMAP and BLAT (Wu & Watanabe 2005; Kent 2002) as aligners, a minimum alignment percent of 75 and a minimum average percent ID of 95. We used *exonerate* (Slater & Birney 2005) to align proteins from nine published plant genomes acquired from Phytozome (Goodstein et al. 2012; *Amaranthus hypochondriacus* v2.1, *Arabidopsis thaliana* v11, *Daucus carota* v2.0, *Helianthus anuus* v1.2, *Lactuca sativa* v5, *Mimulus guttatus* v1.0, *Olea europaea* v1.0, *Solanum lycopersicum* v3.2 and *Solanum tuberosum* v4.03). For *ab initio* predictions, we used four different algorithms. Three of these – *Augustus* (version 3.3.3; Stanke et al. 2006), *GlimmerHMM* (version 3.0.4; Majoros et al. 2004), and SNAP (version 2006-07-28; Korf 2004) – required training to optimise the algorithm for *S. uniflora* and the other – *GeneMark-ES* (version 4.65; Ter-Hovhannisyan et al. 2008) – uses unsupervised training based on the input data. *Augustus* was trained using *BUSCO* (Simão et al. 2015) with the Eudicots gene set. For *GlimmerHMM* and *SNAP*, we used an RNA-seq-based training set to optimise the algorithms. The *pasa_asmbls_to_training_set.dbi* script from *PASA* was first used to produce an initial set of training genes from the *PASA* output which were further filtered to remove single exon, incomplete or invalid gene models using the *prepare_golden_genes_for_predictors.pl* script from the *Just Annotate My Genome* pipeline (Papanicolaou 2014). We used Evidence Modeller (v1.1.1; Haas et al. 2008) to compute weighted consensus gene structure annotations (with evidence weighting of RNA-seq = 10, protein alignment = 5, Augustus annotations = 2 and other *ab initio* annotations = 1, as recommended by the authors) before using PASA to update the EVM consensus predictions and add untranslated regions (UTR) annotations over three iterations. We used GffRead (v0.12.8; Pertea & Pertea 2020) to extract protein sequences and identify genes with non-canonical splice sites, in-frame stop codons, or those missing a start codon or terminal stop codon.

**Supplementary note 6: SeqSNP data processing**.
Trimming was performed using the NGSQCToolKit v2.3.3 (Patel & Jain 2012) IlluQC.pl script with options -l 70 s- 20, followed by Trimmomatic v0.39 (Bolger et al. 2014) using options SLIDINGWINDOW:4:15 and MINLEN:70. Paired reads were mapped to the genome using bwa mem v0.7.17-r1188 (Li 2011), removing PCR duplicates and those with MAPQ<20 using samtools v1.9 (Danecek et al. 2021). Sites were genotyped using bcftools v1.16 (Danecek et al. 2021; Li 2011) sites with QUAL < 15 or DP < 7 were set to missing. Genotypes were merged across the SeqSNP data, and sites with biallelic SNPs, less than 5% missing data, a minor allele frequency greater than 0.2 and a HWE score of > 0 were retained. Following this, individuals with more than 5% missing data across these sites were removed.

**Supplementary note 7. Linkage map construction.**
We used a pedigree constructed using two dummy parents. The SeparateChromosomes2 function was used to assign markers to chromosomes with parameter distortionLod=1 and minimum linkage group size 25 markers: LOD values between 1 and 55 were tested, and the LOD value was chosen (13) based on the highest number of markers assigned to linkage groups, and the number of linkage groups equalling the expected number of chromosomes (12). Additional unassigned

markers were added to these linkage groups using the function JoinSingles2All with parameters lodLimit=10, lodDifference=8, distortionLod=1 and iterate=1. Marker order within each linkage group was assessed using the OrderMarkers2 function, with parameters interference1=1 interference2=2 sexAveraged=1. This was run 20 times for each linkage group, the initial marker order was set by preserving the order of markers within the scaffolds, but randomly assigning the orientation of these scaffolds and the order of scaffolds within the linkage group. Following the 20 runs, the order with the highest log-likelihood was chosen. The resulting chromosome maps were trimmed using the LepWrapTrim.R function of LepWrap (Dimens 2022), removing markers from the end 20% of maps that had gaps of >2cM from the rest of the map.

**Supplementary note 8. Synteny analysis.**
Orthologs between *S. uniflora* and the other species were first identified using the *ortholog* command in the jcvi.compara.catalog module using a C-score cutoff of 0.99. We used *screen* command from the jcvi.compara.synteny module (with options --minspan=30 –simple) to identify syntenic blocks. Synteny plots were created using the jcvi.graphics.karyotype module.

**Supplementary figures**

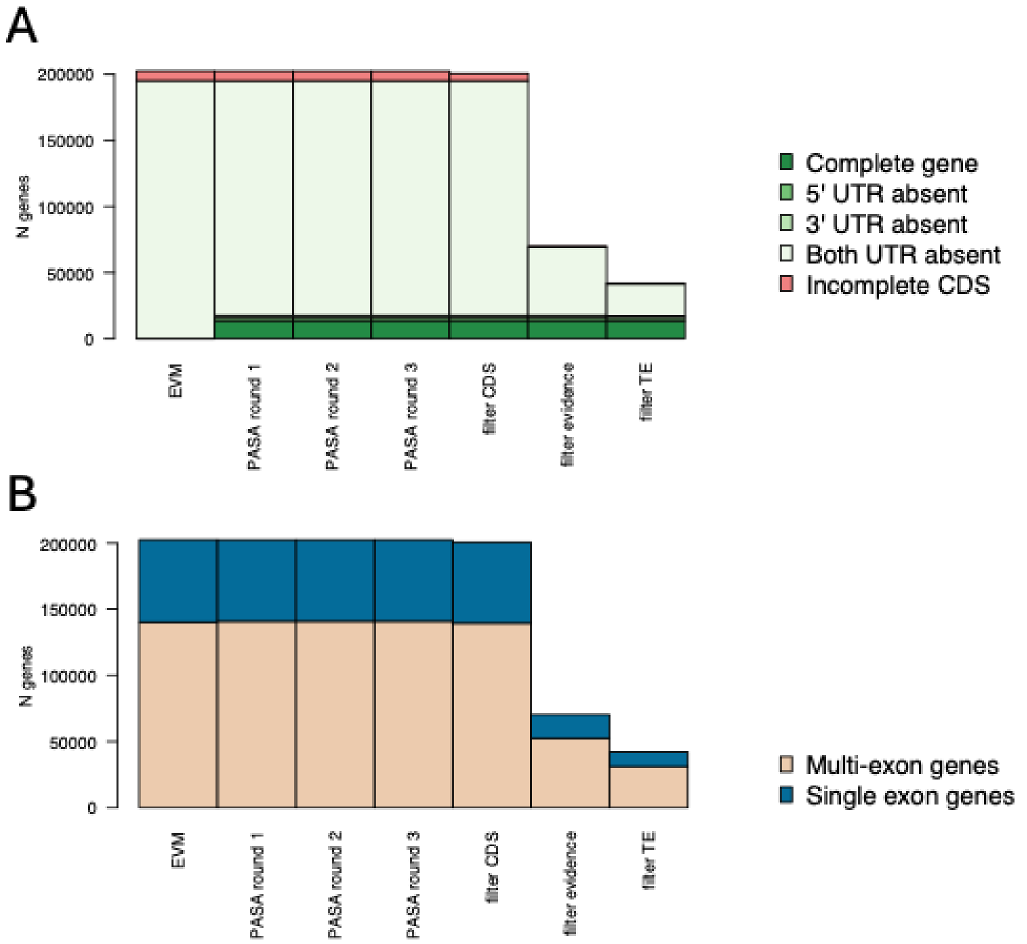


Fig S1. Bar plots show the number of genes at each level of completeness (A) and complexity (B) at each stage of annotation consolidation and filtering.

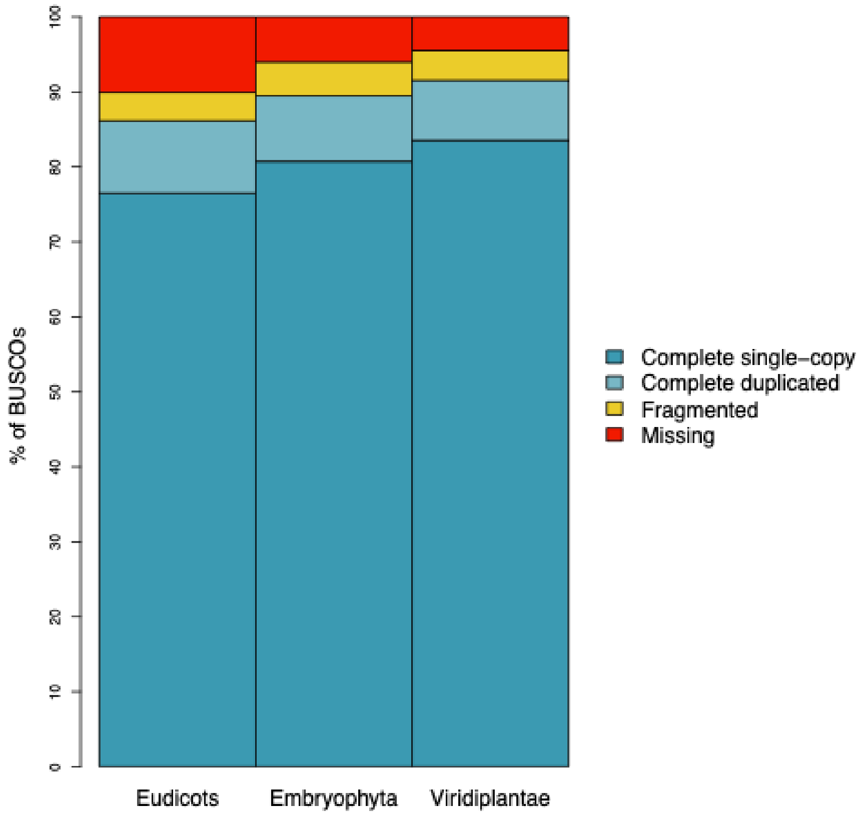


Fig S2. Completeness of the final annotation as estimated by BUSCO, using three BUSCO reference datasets. Bar plots show the proportion of BUSCO single copy orthologs in each BUSCO category.

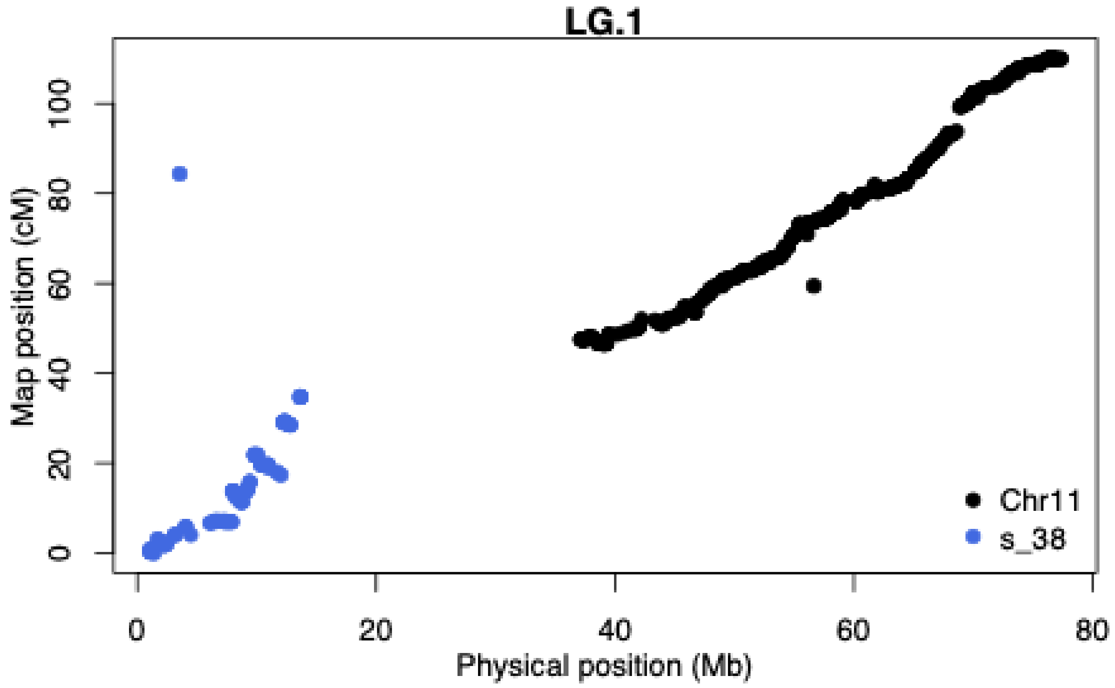


Fig. S3. Estimated placement of scaffold > 1Mb on linkage group (LG) 1. Physical position versus map position is shown for all markers. For each scaffold, starting physical position is estimated from it's starting map position scaled by the LG-wide recombination rate.

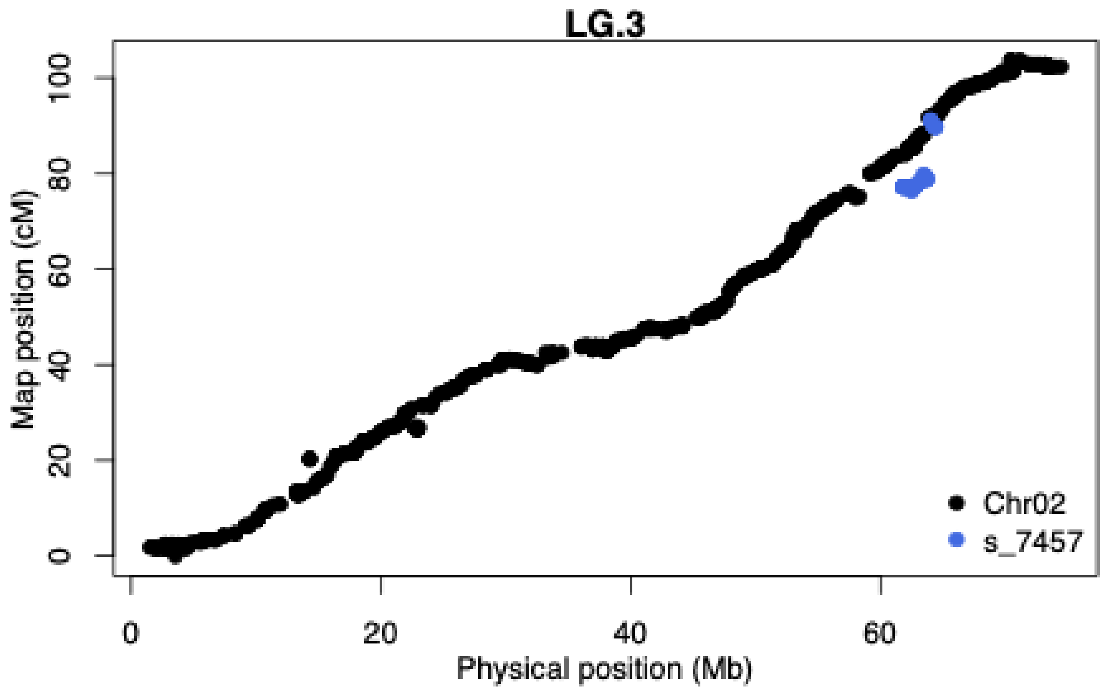


Fig. S4. Estimated placement of scaffold > 1Mb on LG 3. Formatting as in Fig. S3

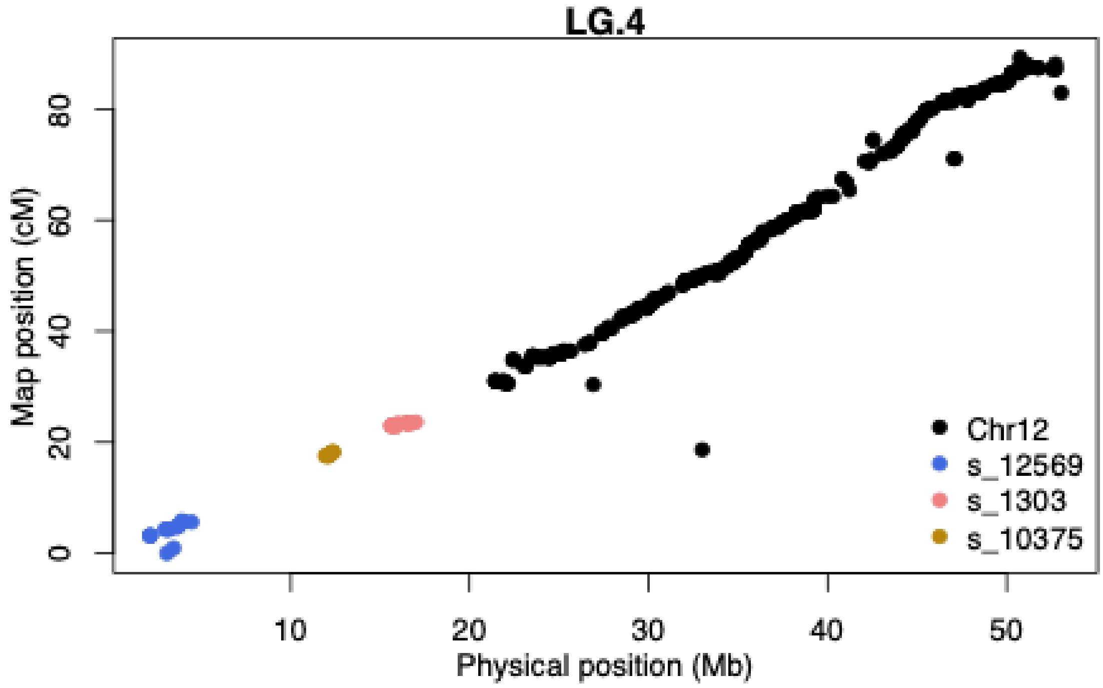


Fig. S5. Estimated placement of scaffold > 1Mb on LG 4. Formatting as in Fig. S3.

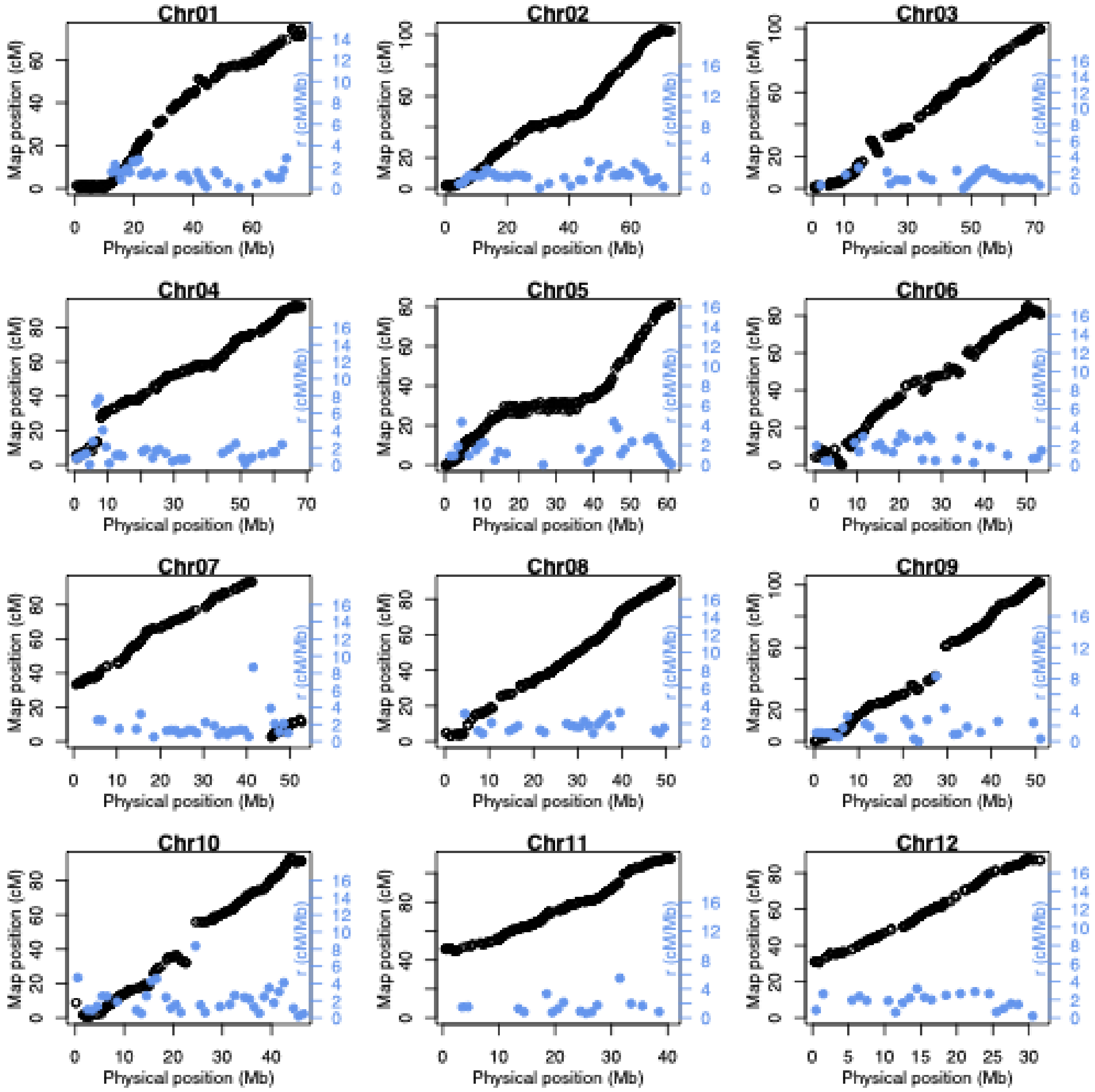


Fig. S6. Local recombination rate estimation for all chromosomes. Black points show the relationship between physical position (Mb; x-axis) and linkage map position (cM; left y-axis). Blue points show local recombination rate (cM/Mb; right y-axis) estimated in non-overlapping 1Mb windows across each chromosome. Outlier markers were not used for recombination rate estimation and are not shown.

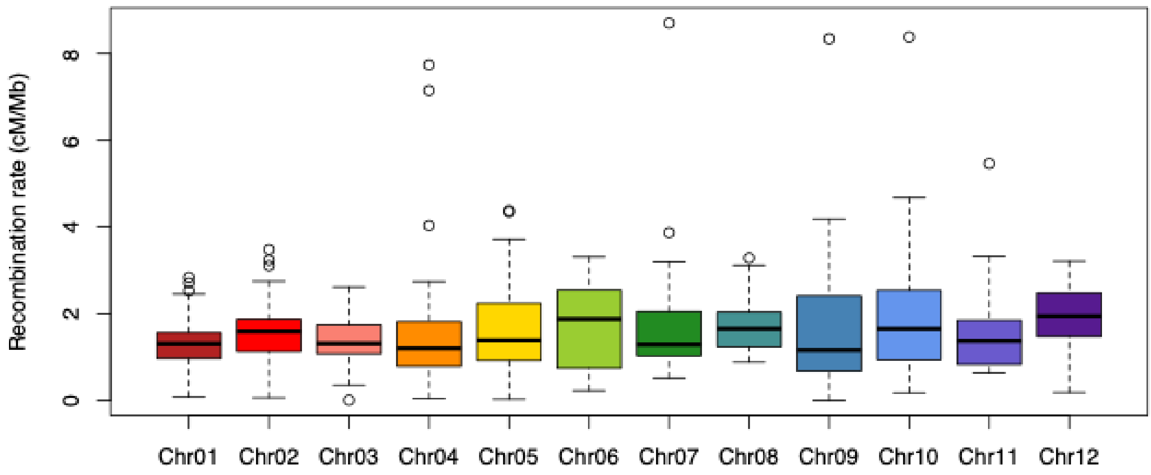


Fig. S7. The distribution of local recombination rate for each chromosome. Each boxplot shows the distribution of recombination rate for a single chromosome (coloured as in Fig. 1). Outlier windows are shown as black circles on each boxplot.

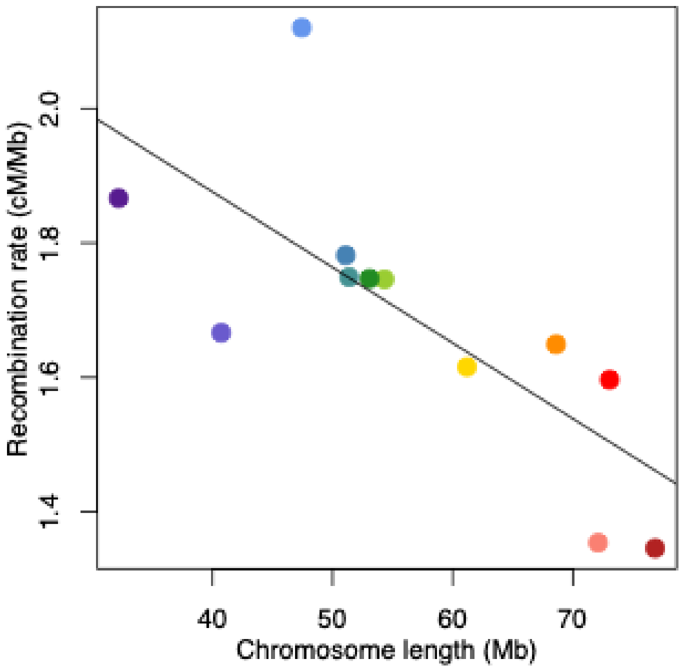


Fig. S8: the relationship between chromosome length and mean local recombination rate. Each point represents one chromosome (coloured as in Fig. 1). The line shows a linear regression line. The negative correlation was strong and significant (Spearman's rank correlation: $\rho$ = -0.87; *P* = 0.0004).

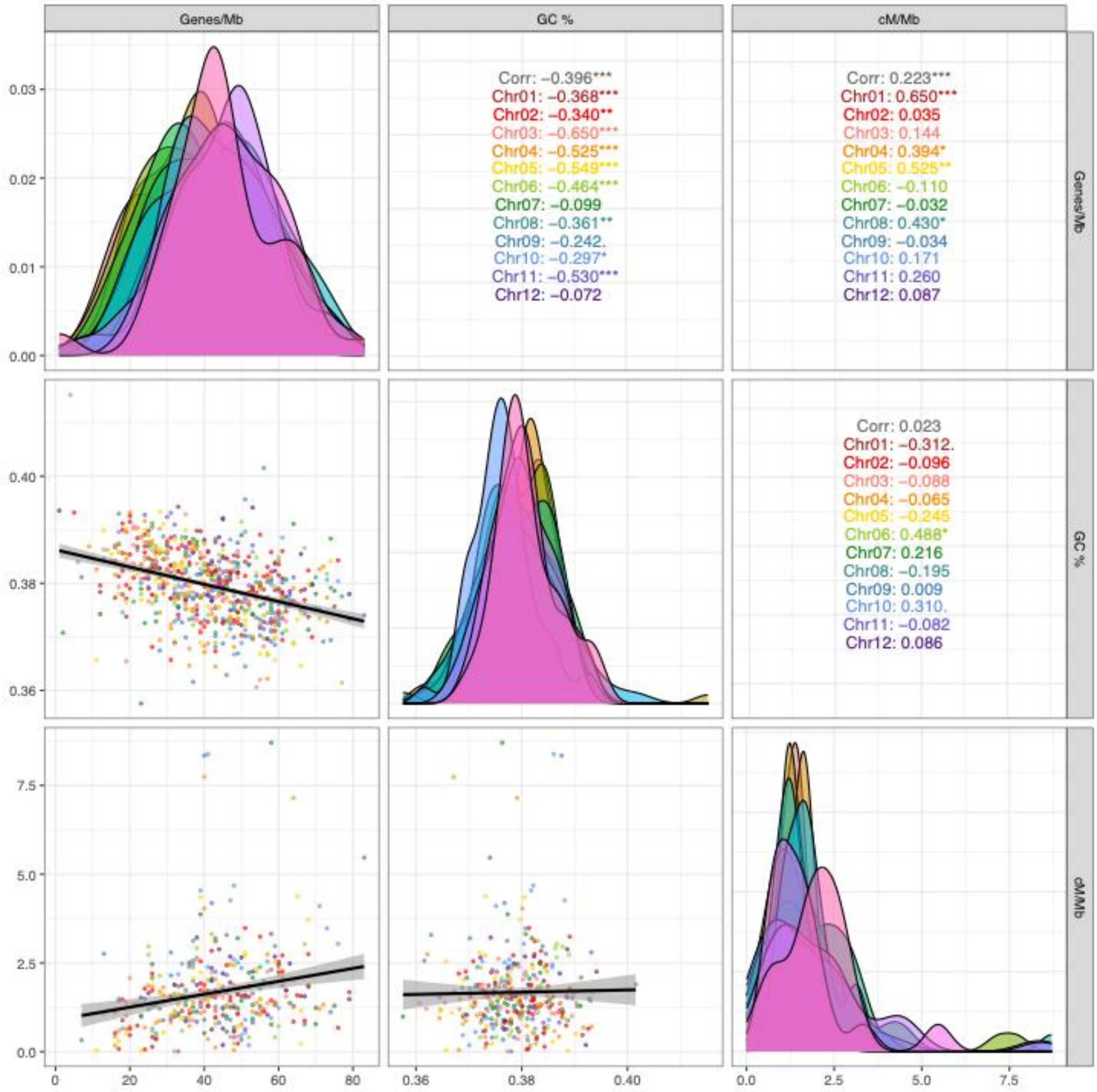


Fig. S9. The relationship between gene density (genes per 1 Mb), GC proportion and local recombination rate (cM/Mb). Text in the upper panels shows the Spearman's correlation for the whole dataset and each chromosome individually. Significance is indicated by stars ($P$ < *0.1* = "."; $P$ < 0.05 = "*"; $P$ < 0.01 = "**"; $P$ < 0.001 = "***"). Density plots in the diagonal panels show the distribution of each statistic separately for each chromosome (coloured as in upper panels). Dot plots in lower panels show the relationship between with each statistic. Points are coloured by chromosome (as in upper panels) and a trendline with 95% confidence intervals is shown.
